# Activation of Infralimbic cortex neurons projecting to the nucleus accumbens shell suppresses discriminative stimulus-triggered relapse to cocaine seeking in rats

**DOI:** 10.1101/2025.06.23.661090

**Authors:** Hajer E Algallal, Isabel Laplante, Domiziana Casale, Shaghayegh Najafipashaki, Audrey Pomerleau, Thomas Paquette, Anne-Noël Samaha

## Abstract

Cocaine addiction is marked by high relapse rates, often triggered by drug-associated cues. These cues can be conditioned stimuli (CSs), which occur after drug intake and are paired with drug effects, and discriminative stimuli (DSs), which signal drug availability, regardless of ongoing drug-seeking behaviour. While projections from the infralimbic cortex (IL) to the nucleus accumbens (NAc) shell are known to regulate CS-induced cocaine relapse, their role in DS-triggered relapse is not known. To investigate this, we examined how activating IL→NAc shell projections influences relapse driven by DSs and CSs during abstinence from intermittent cocaine use. Female Sprague-Dawley rats received viral-mediated gene expression of excitatory designer receptors exclusively activated by designer drugs in the IL. Rats then self-administered cocaine during 12 intermittent-access sessions (5-min cocaine ON/25-min cocaine OFF, 4h/day). A discrete light (DS+) signalled drug-available periods, while a different light (DS-) signalled drug non-availability. During each DS+ period, cocaine infusions were paired with a compound light-tone (CS+). Four weeks later, rats were tested for cue-induced cocaine seeking following response-independent presentation of DS+, CS+ or both. Immediately prior to testing, rats received intra-NAc shell clozapine N-oxide or aCSF to activate IL terminals. DS+ alone and DS+/CS+ combined triggered greater cocaine seeking than did the CS+. Activation of IL→NAc shell projections suppressed relapse behaviour in DS+ and DS+/CS+ conditions. These findings highlight the distinct and powerful influence of DSs on relapse and identify the IL→NAc shell circuit as a promising target for relapse prevention.

## Introduction

Substance use disorders are a leading cause of psychopathology and death in the world [1]. A major challenge for treatment is the high rate of relapse, where most people who stop using drugs relapse within a year [2]. Cocaine users often cite *people*, *places* and *things* associated with drug use as triggers for drug cravings [3] and laboratory studies confirm that such cues trigger craving and relapse in drug users [4, 5]. Thus, understanding how drug cues influence brain and behaviour is essential for addressing relapse.

Animal models are widely used to study cue-induced relapse [6, 7] and they typically involve relapse-like behaviour induced by drug-paired conditioned stimuli (CSs) [8–13]. For instance, during training, rats press a lever for drug infusions and CS presentation (e.g., a light/tone). After abstinence, a test measures the extent to which the CS reinforces lever pressing, without drug, as a measure of relapse to drug seeking. During both training and testing, the CS is contingent on the drug-seeking response. That is, the CS and the neural substrates it recruits operate *after* drug seeking is initiated. Relapse to drug use also involves stimuli that signal drug availability and act as *triggers* of drug-seeking actions, before any such actions have been initiated [14–17]. These discriminative stimuli (DS) are unique in that they evoke a conditioned motivational state that can prompt drug craving and relapse before an individual is even aware of wanting drugs [18, 19].

Neural circuits involved in DS-triggered relapse have seldom been studied [20–22]. Through cortico-limbic-striatal circuits, the medial prefrontal cortex (mPFC) contributes to the execution and inhibition of reward-seeking behaviours [23–27]. Within the mPFC, neuronal activity in the infralimbic cortex (IL) promotes reward seeking in the presence of a DS signalling reward availability (DS+) and inhibits responding in the presence of a DS signalling reward unavailability (DS-) [28–35]. However, the IL-dependent neural pathways involved are unknown. A possibility is the glutamatergic pathway from the IL to the nucleus accumbens (NAc) shell [36–39]. Notably, under conditions where cocaine-seeking responses are reinforced by a previously drug-paired CS—that is, when the drug cue is encountered *after* a drug-seeking response—activating IL-to-nucleus accumbens (NAc) shell neurons suppresses cocaine seeking [40, 41].

This raises the question; can activity in IL-to-nucleus accumbens shell neurons suppress relapse triggered by drug cues *before* any drug-seeking behaviour begins? To address this, we used viral-mediated gene transfer of designer receptors (DREADDs) to activate these neurons and determine effects on cocaine relapse-like behaviour triggered by a DS and CS when these cues are presented independent of behaviour.

## Materials and methods

Supplementary Methods provide details on experimental subjects, drugs, apparatus, procedures and statistical analyses. All experiments followed the Canadian Council of Animal Care and were approved by the animal care committee of the Université de Montréal.

### Experiment 1: Effects of chemogenetic activation of IL**→**NAc Shell projections on cocaine seeking induced by discriminative and conditioned stimuli

Here we determined in female rats the extent to which increased activity in IL→NAc Shell neurons suppresses DS- and CS-induced cocaine seeking when these cues are presented response-independently during abstinence. Fig. 1 (top) shows the sequence of experimental events. Before any cocaine self-administration experience, we first determined the Pavlovian conditioned approach phenotype of our rats, because goal-trackers are reported to respond more to DSs and sign-trackers more to CSs [42–47], cf. [48–50]. We then trained the rats to press a lever to self-administer sucrose, before learning to lever press for cocaine (0.5 mg/kg/inf, injected over 5 s). Cocaine was available under intermittent-access (IntA) conditions (Fig. 2B), during which cocaine was available under FR3 during 5-min ON periods signalled by a DS+ (cue light above the left lever) and then unavailable during 25-min OFF periods signalled by a DS-(cue light on the back wall). During DS+ periods, each cocaine infusion was also paired with a 5-s CS+ (light+tone cue). After IntA sessions, 36 rats received a viral vector containing a constitutive G_q_-coupled designer receptor exclusively activated by a designer drug (AAV8-CaMKIIα-hM3D(G_q_)-mCherry; Canadian Neurophotonics Platform - Viral Vector Core, QC, Canada) and 20 rats received a viral vector containing mCherry alone (AAV8-CaMKIIα-mCherry; Canadian Neurophotonics Platform - Viral Vector Core, QC, Canada). All rats also had bilateral stainless steel guide cannula (23 gauge; HRS Scientific, Anjou, QC, Canada) implanted into the NAc Shell. Following 6 weeks of forced abstinence (and 5-6 weeks after adeno-associated virus microinjections) during which rats stayed in their home cages, we assessed cocaine-seeking behaviour (non-reinforced lever pressing) induced by response-independent presentation of the DS+, CS+, DS-and DS+/CS+ combined (three, 2-min presentations/cue; pseudorandomized order). Rats received bilateral CNO (1mM; 0.5 μl/hemisphere) or aCSF microinfusions into the NAc shell ≤ 10 min before the test (between-subjects).

**Figure 1.**
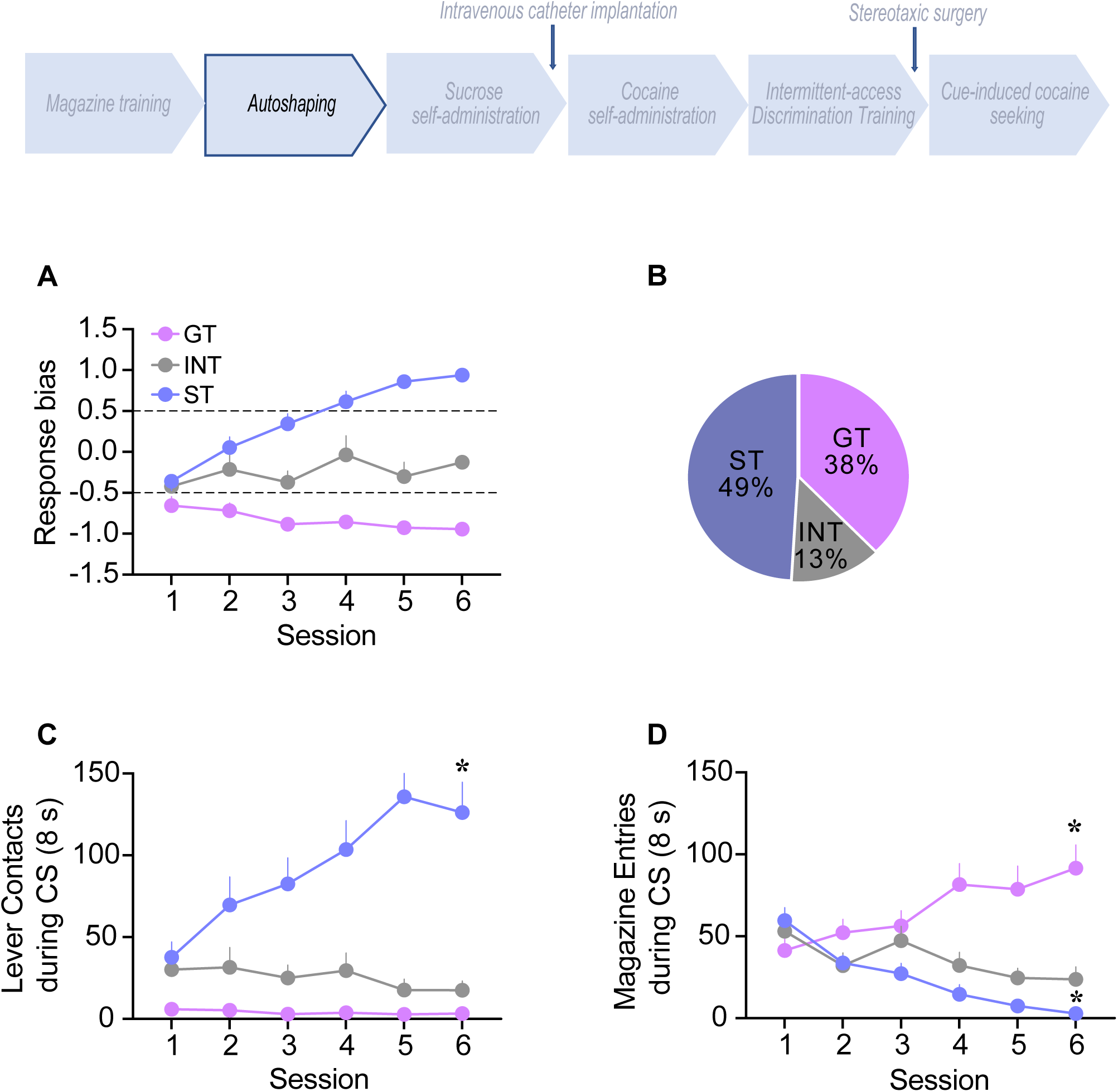
Acquisition of Pavlovian conditioned responding across autoshaping sessions. **A** Pavlovian conditioned response bias scores across autoshaping sessions for goal-tracker rats (GT; scores between −0.5 and −1.0), sign-tracker rats (ST; scores between +0.5 and +1.0) and intermediates (INT; scores between −0.49 and +0.49). **B** The percentage of rats with each Pavlovian conditioned response phenotype. **C** Lever contacts during presentation of the lever conditioned stimulus (CS), and **D** Magazine entries during presentation of the lever CS. * *p* < 0.001, session 6 vs. session 1. Data are mean ± SEM, n = 56.

**Figure 2.**
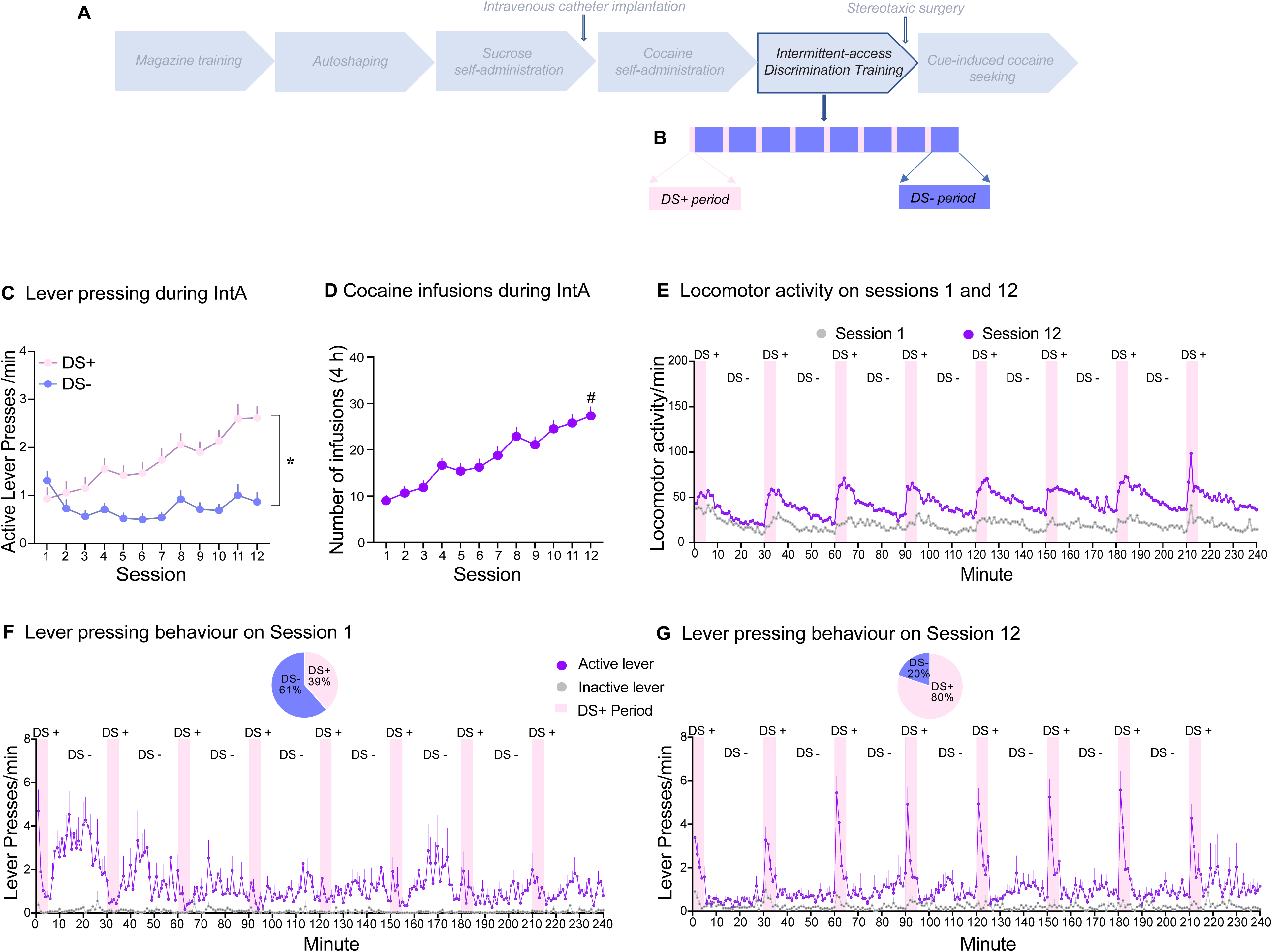
Intermittent-access cocaine self-administration (IntA) under the control of discriminative stimuli signaling cocaine availability (DS+) and unavailability (DS-). **A** The experimental timeline. **B** Schematic of a 4-h IntA session during which 5-min cocaine ON periods signaled by a DS+ cycled with 25-minute cocaine OFF periods signaled by a DS-. **C** Across sessions, rats learned to increase rates of active lever pressing during cocaine ON/DS+ periods and suppress responding during cocaine OFF/DS-periods. **D** Rats escalated their drug intake across IntA sessions. **E** By Session 12, locomotor activity peaked during cocaine ON/DS+ periods and abated during cocaine OFF/DS-periods, and locomotion rates were also greater than on Session 1. **F** On Session 1, lever pressing rates were higher during DS-vs. DS+ periods, even though cocaine was only available during DS+ periods. The inset shows that rats only made 39% of their total active lever presses during DS+ periods, and 61% during DS-periods. **G** By Session 12, rates of active lever pressing peaked during DS+ periods and responding was suppressed during DS-periods. The inset shows that rats now made 80% of their active lever presses during DS+ periods. * *p* < 0.001, DS+ vs. DS-, Session 12. # *p* < 0.001, Session 12 vs Session 1. Data are mean ± SEM, n = 56.

Because human cocaine users also encounter drug-associated cues while experiencing the pharmacological effects of cocaine, a secondary goal was to determine how being on cocaine might influence cue-induced cocaine-seeking behaviour. Two days after the first cue-induced cocaine-seeking test, rats were given a second cue-induced cocaine-seeking test, with no brain microinjections. Instead, the rats received cocaine (10 mg/kg, i.p.) or saline (i.p.) ≤ 5 min prior to the test (between-subject). Two days after this 2^nd^ test, we validated G_q_ DREADD functionality by quantifying the effects of chemogenetic activation of IL→NAc shell neurons on c-Fos protein expression in the NAc shell. Half the rats expressing G_q_ DREADD-mCherry and half expressing mCherry alone received bilateral microinjections of 1 mM CNO into the NAc shell, and the other half received aCSF. Ninety minutes later, brains were extracted and processed for histology and c-Fos immunohistochemistry.

### Experiment 2: Effects of intra-NAc shell CNO injections on spontaneous locomotion

Here we determined the extent to which chemogenetic activation of IL→NAc shell neurons influenced general motor activity, to consider potentially non-specific effects. Thus, 4-5 weeks after adeno-associated virus microinfusions, all rats received a test for spontaneous locomotor behaviour. Approximately 10 min before testing, half the rats expressing G_q_ DREADD-mCherry or mCherry alone received bilateral microinjections of 1 mM CNO into the NAc shell, and the other half received aCSF. The rats were then placed in locomotor activity cages and horizontal locomotion was recorded for 1 h. Ninety min after the microinjections, rats were perfused, and brains were extracted for histology and c-Fos immunohistochemistry as in Exp. 1.

## Results

### Experiment 1: Effects of chemogenetic activation of IL→NAc Shell neurons on cocaine seeking induced by discriminative and conditioned stimuli

#### Autoshaping

Fig. 1A shows the Pavlovian conditioned response bias score across autoshaping sessions in sign-trackers, goal-trackers and intermediates. By the end of autoshaping, 49% of the rats were sign-trackers, 38% were goal-trackers, and 13% were intermediates (Fig. 1B). During presentation of the lever CS, sign-trackers increased their lever contacts across sessions (Fig. 1C; Session 1 vs. 6, *z* = −4.423, *p* < 0.001) and decreased their magazine entries (Fig. 1D; Session 1 vs. Session 6, *z* = −4.70, *p* < 0.001). In contrast, goal-trackers maintained low and stable rates of lever contacts (Fig. 1C; *p* > 0.05) and increased their magazine entries (Fig. 1D; Session 1 vs. Session 6, *z* = −3.25, *p* = 0.001). Thus, all rats learned a conditioned response, such that for sign-trackers and goal-trackers, CS-reward pairings increased the rate of approach during CS presentation to respectively, the cue or the goal [51].

### Cocaine Self-Administration Under the Control of Discriminative Stimuli

Fig. 2 By IntA Session 12, rats pressed more on the active lever during DS+ vs. DS-periods (Fig. 2C; Session 12, DS+ vs. DS-, *z* = −5.77, *p* < 0.001). Rats also escalated both their cocaine intake (Fig. 2D; Session 1 vs. 12, *z* = −5.77, *p* < 0.001) and their locomotor activity (Fig. 2E; *z* = −5.16, *p* < 0.001) from Session 1 to 12. Even though cocaine was only available during DS+ periods, rates of lever pressing behaviour during Session 1 were higher during DS-periods (Fig. 2F), such that rats made 61% of their total active lever presses during DS-periods and only 39% during DS+ periods (Fig. 2F inset). However, on Session 12 (Fig. 2G), responding now peaked during DS+ periods, such that rats made most (80%) of their active lever presses during DS+ and only 20% during DS-periods (Fig. 2G inset). Thus, cocaine self-administration came under control of the DSs, with rats lever pressing for cocaine in the presence of the DS+ and suppressing responding in the presence of the DS-.

### Cue-induced cocaine seeking following intra-NAc Shell aCSF and CNO injections

Following IntA discrimination training, we assessed cocaine-seeking behaviour (with no cocaine available) in response to the DS+, DS-, CS+ and DS+CS+ combined, during a test where each cue type was presented response-independently on 3 trials, in pseudorandom order. Half of the mCherry-expressing and the G_q_ DREADD-expressing rats received intra-NAc Shell CNO, the other half received aCSF. The three control groups (mCherry-aCSF, mCherry-CNO, G_q_ DREADD-aCSF) showed similar lever-pressing behaviour during the test and were pooled under ‘Control’. Control rats pressed significantly more on the active lever during presentation of the DS+ and DS+CS+ combined relative to all other conditions (Fig. 3A; DS+ vs. inter-trial interval or ITI, *z =* −3.83*, p <* 0.001; DS+ vs. CS+, z = −4.17, *p <* 0.001; DS+ vs. DS-, z = −4.19, *p <* 0.001; DS+CS+ vs. ITI, z = −3.57, *p <* 0.001; DS+CS+ vs. CS+, *z =* −3.96, *p <* 0.001; DS+CS+ vs. DS-, *z =* −3.67*, p <* 0.001). In contrast, responding to the CS+ was similar to that during the ITI (z = - 0.80, *p >* 0.05). Thus, relative to when no cue was presented (i.e., during the ITI), only the DS+presented alone or in combination with the CS+ triggered significant increases in cocaine-seeking responses. In addition, the DS+ and DS+CS+ combined prompted higher rates of cocaine-seeking behaviour compared to the CS+. Thus, the DS+ triggered peaks in cocaine-seeking responses, while the CS+ was ineffective.

**Figure 3.**
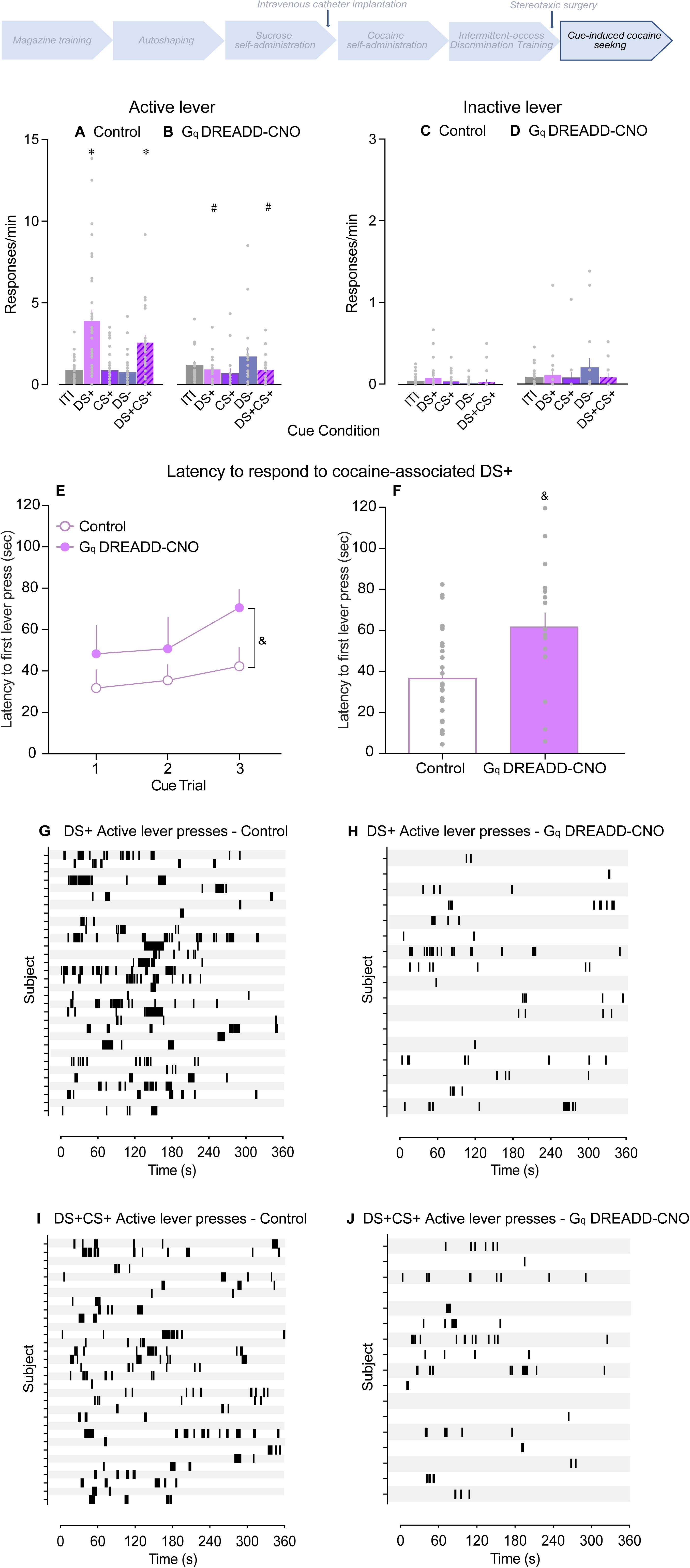
Chemogenetic activation of IL→NAc Shell neurons suppresses the increases in cocaine seeking triggered by a drug-predictive discriminative stimulus. Six weeks after forced abstinence from intermittent-access cocaine self-administration, rats received a cue-induced cocaine-seeking test during which no cocaine was available. Pressing on the previously cocaine-associated (active) lever following response-independent presentation of the DS+, DS-, CS+ and DS+/CS+ combined or during the inter-trial interval (ITI) in **A** Control rats and **B** rats with chemogenetic activation of G_q_ DREADD-expressing terminals from the infralimbic cortex to the nucleus accumbens shell (G_q_ DREADD-CNO rats). Inactive lever presses per min in **C** Control rats and **D** G_q_-DREADD-CNO rats. All cues were presented independent of responding and lever pressing produced no outcome. Latency to initiate the first active lever press **E** following each of the three DS+ trials of the session in Control and G_q_ DREADD-CNO rats and **F** averaged across the three DS+ trials in each group. Timestamps of active lever presses in individual subjects during DS+ presentation in **G** Control and **H** G_q_ DREADD-CNO rats, and during presentation of the DS+CS+ combined in **I** Control and **J** G_q_ DREADD-CNO rats. * *p* < 0.0001, vs ITI, CS+ & DS-. # *p* < 0.005, DS+ & DS+CS+ in Control rats vs DS+ & DS+CS+ in G_q_ DREADD-CNO rats. & *p* < 0.05 G_q_ DREADD-CNO vs Control. Data are mean ± SEM, Control n = 32, G_q_ DREADD-CNO n = 17.

Compared to Control rats (Fig. 3A), G_q_ DREADD-CNO rats responded significantly less to the DS+ (Fig. 3B; *z =* −3.18, *p =* 0.001) and DS+CS+ combined (Fig. 3B; *z =* −2.98, *p =* 0.003). In G_q_ DREADD-CNO rats, CNO had no effect on responding during presentation of the CS+ (z = −0.42, *p* = 0.75), DS-(z = −0.98, *p* = 0.33) or ITI (z = −0.58, *p* = 0.56), or on inactive lever pressing (Fig. 3D vs. 3C; DS+, z = −0.46, *p =* 0.64; DS+/CS+, z = −1.79, *p =* 0.07; CS+, z = −0.61, *p <* 0.54; DS-, z = −0.37, *p <* 0.71; ITI, z = −0.89, *p <* 0.37). Fig 3E shows the latency to initiate active lever pressing upon DS+ presentation as a function of trial number. Compared to Control rats, G_q_ DREADD-CNO rats waited longer to initiate responding, both when we analysed latency as a function of the three DS+ trials of the session (Fig 3E; Group effect, F_1,43_= 6.99, *p* = 0.01; no other comparisons were statistically significant) and averaged across trials (Fig 3F; Control v. G_q_ DREADD-CNO, *p* = 0.003). There were no group differences in the latency to respond to DS+CS+ presentation as a function of the three DS+CS+ trials of the session (data not shown; Group effect, F_1,42_= 0.09, *p* = 0.76) or averaged across trials (data not shown; Control v. G_q_ DREADD-CNO, *p* > 0.05). To further highlight the suppression in responding in G_q_ DREADD-CNO rats, Figs. 3G-H show active lever pressing records during DS+ presentation in respectively, individual Control and G_q_ DREADD-CNO rats, and Figs. 3I-J show responding during DS+CS+ presentation in the two groups. Thus, increasing activity in IL→Shell neurons significantly suppressed the increases in cocaine-seeking behaviour normally triggered by a DS+ by reducing both the rate and latency of responding.

### Cue-induced cocaine seeking while under the effects of cocaine

Two days following the cue-induced cocaine-seeking test in Fig. 3, we gave rats a second cue-induced cocaine-seeking test to assess the effects of an i.p. cocaine (or saline) injection on responding. Under saline, rats pressed more on the active lever during the DS+ vs. the ITI (Fig. 4A; *z =* −3.11*, p <* 0.002), but responding no longer differed across cue types (Fig. 4A; DS+ vs. CS+, z = −1.85, *p <* 0.07; DS+ vs. DS-, z = -0.17, *p =* 0.87; DS+CS+ vs. ITI, z = -1.63, *p <* 0.10; DS+CS+ vs. CS+, *z =* -1.26, *p =* 0.21; DS+CS+ vs. DS-, *z =* -0.96*, p <* 0.34; CS+ vs. ITI, z = - 1.66, *p =* 0.10; DS-vs. ITI, *z =* -1.98, *p =* 0.06). This suggests that (DS+)-induced increases in cocaine-seeking behaviour extinguished with repeated testing and/or the passage of time (also compare Figs. 3A and 4A). A cocaine injection reinstated responding to the DS+ and DS+CS+ combined. First, under cocaine, rats again responded most during the DS+ relative to other cue conditions (Fig. 4B; vs. ITI, *z =* -3.78*, p <* 0.001; vs. CS+, z = -4.12, *p <* 0.001; vs. DS-, z = -3.57, *p <* 0.001). They also responded more during the DS+CS+ than they did during CS+ presentation (*z =* -2.62, *p =* 0.009) or DS-presentation (*z =* -2.07*, p =* 0.04). The effects of cocaine were specific to the DS+ and DS+CS+ conditions; cocaine did not increase responding to either the CS+ (z = - 1.95, *p* = 0.05) or the DS-(z = -1.53, *p* = 0.13) beyond that seen during the ITI. Second, compared to saline (Fig. 4A), cocaine significantly increased rates of active lever pressing during presentation of the DS+ (Fig. 4B; *z =* -3.09, *p =* 0.002), DS+CS+ combined (*z =* -2.54, *p =* 0.01) and ITI (*z =* -2.53, *p =* 0.01). In contrast, relative to saline, there was no effect of cocaine on responding to the CS+ (*z =* -1.67, *p =* 0.10) or DS-(*z =* -0.65, *p =* 0.52). Cocaine also had no effects on inactive lever pressing (Fig. 4D; DS+, z = -1.59, *p =* 0.11; DS+CS+, z = -1.88, *p =* 0.06; DS-, z = -0.39, *p <* 0.70; ITI, z = -1.77, *p <* 0.08), except for an increase in responding during CS+ presentation (Fig. 4A*; z =* -2.46, *p =* 0.01). Fig 3E shows the latency to initiate active lever pressing upon DS+ presentation as a function of trial number. Relative to saline, cocaine did not significant influence the latency to respond to the DS+ (Fig 3E; Group effect, F_1,74_= 3.31, *p* = 0.07; no other comparisons were statistically significant; Fig 3F; *p* > 0.05). Similarly, there were no group differences in the latency to respond to the DS+CS+ combined as a function of the three DS+CS+ trials of the session (data not shown; Group effect, F_1,38_= 0.36, *p* = 0.55) or averaged across trials (data not shown; *p* > 0.05). To further highlight the stimulatory effects of cocaine on (DS+)-induced drug seeking, Figs. 4G-H show active lever pressing records during DS+ presentation in respectively, individual Control and G_q_ DREADD-CNO rats, and Figs. 4I-J show responding during DS+CS+ presentation in the two groups. Thus, re-exposure to cocaine reinstated (DS+)-induced cocaine-seeking behaviour by increasing rates of responding, without influencing the latency to respond.

**Figure 4.**
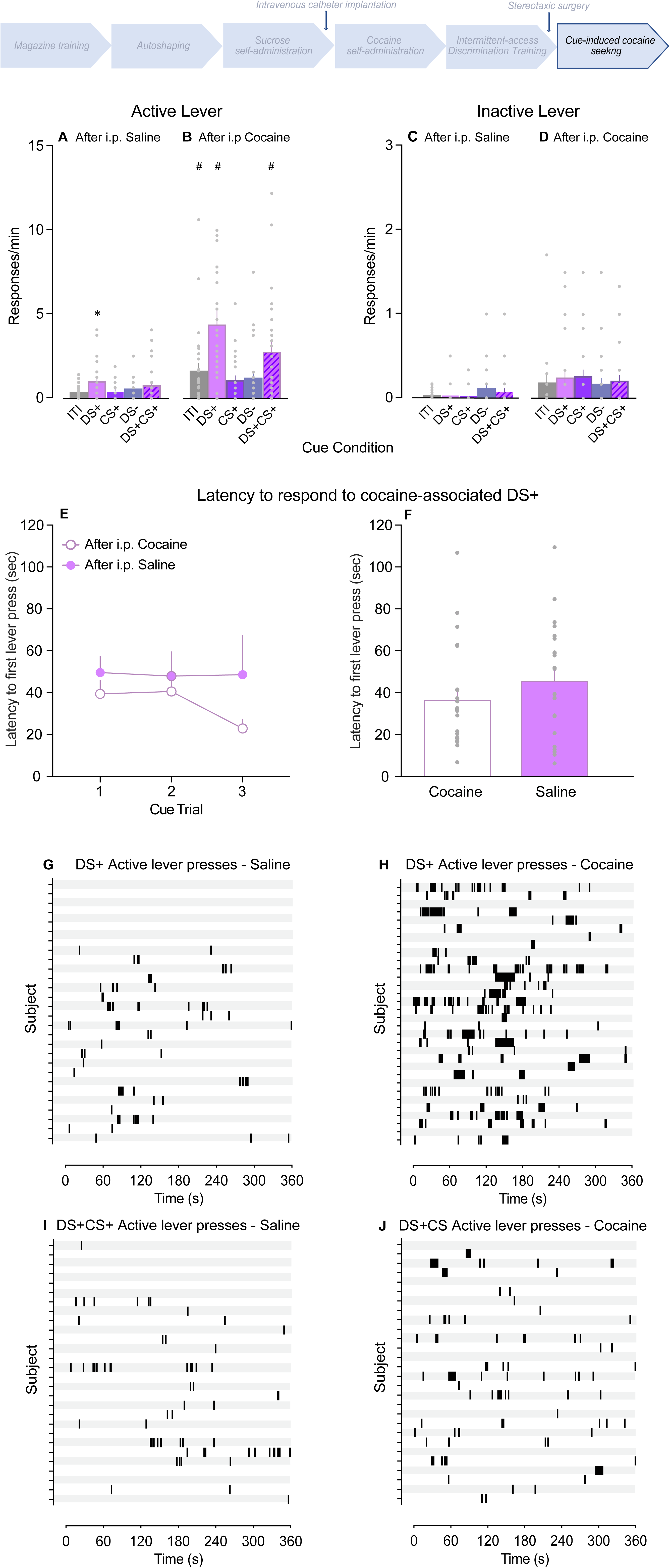
Re-exposure to cocaine reinstates cocaine seeking triggered by presentation of a drug-predictive discriminative stimulus. Six weeks after forced abstinence from intermittent-access cocaine self-administration, rats received a cue-induced cocaine-seeking test during which no cocaine was available. Immediately before the test, rats received an i.p. injection of cocaine or saline. Pressing on the previously cocaine-associated (active) lever following response-independent presentation of the DS+, CS+, DS- and DS+/CS+ combined or during the inter-trial interval (ITI) in **A** Saline-injected rats and **B** Cocaine-injected rats. Inactive lever presses per min in **C** Saline-injected rats and **D** Cocaine-injected rats. All cues were presented independent of responding and lever pressing produced no outcome. Latency to initiate the first active lever press **E** following each of the three DS+ trials of the session in cocaine-vs. saline-injected rats and **F** averaged across the three DS+ trials in each group. Timestamps of active lever presses in individual subjects during DS+ presentation in **G** Saline-injected and **H** Cocaine-injected rats, and during presentation of the DS+CS+ combined in **I** Saline-injected and **J** Cocaine-injected rats. * *p <* 0.002, vs. ITI. # *p* ≤ 0.01, DS+, DS+CS+ & ITI Cocaine vs respectively, DS+, DS+CS+ & ITI Saline. Data are mean ± SEM, Saline n = 28, Cocaine n = 28.

### More cocaine taken in the past predicts greater responding to drug-associated cues

We observed individual variability in responding to the cocaine-associated cues. Table 1 shows correlations between cumulative intake across IntA sessions or Pavlovian conditioned response bias score and later cocaine-seeking behaviour in response to the DS+, CS+ and DS+CS+ combined. There were no significant correlations between Pavlovian conditioned response score and lever pressing behaviour during cue presentation (DS+, CS+ or DS+CS+; all *P* values > 0.05). Thus, a goal-vs. sign-tracker phenotype did not predict cocaine-seeking behaviour in the presence of drug-associated DSs or CSs. However, the more cocaine rats took the more they later responded for cocaine during presentation of the DS+, CS+ and DS+CS+ combined (Table 1; all *P* values < 0.05).

### Validation of G_q_ DREADD-mediated neuronal activation (Experiments 1 & 2)

We used c-Fos protein expression as a marker of cell activity to confirm activation of NAc shell cells following CNO-mediated stimulation of G_q_ DREADD-expressing IL inputs. Rats expressing mCherry or G_q_ DREADD-mCherry received intra-NAc shell infusions of aCSF or CNO before c-Fos quantification. In Experiment 1 (Fig. 5E), G_q_ DREADD-mCherry rats receiving intra-NAc shell CNO infusions showed significant increases in c-Fos+ cells in the NAc shell compared to the control groups (vs. mCherry-aCSF, z = - 3.21, *p* < 0.001; vs. mCherry-CNO, z = - 4.07, *p* < 0.001; vs. G_q_ DREADD-aCSF, z = - 4.25, *p* < 0.001). We saw the same effect in Experiment 2 (Fig. 5G; G_q_ DREADD-CNO vs. mCherry-aCSF, z = - 2.13, *p* = 0.03; vs. mCherry-CNO, z = - 2.02, *p* = 0.04; vs. G_q_ DREADD-aCSF, z = - 2.12, *p* = 0.03). Thus, chemogenetic activation of IL cortex neurons significantly increased c-Fos protein expression in the NAc shell. In Exp. 1, we saw some virus expression in the ventral aspect of the prelimbic cortex (Fig. 5C, left). Prelimbic cortex neurons project to the NAc core, and these projections also mediate cue-induced cocaine-seeking behaviour [52–55]. Thus, we also quantified CNO-induced c-Fos+ cells in the NAc core. In G_q_ DREADD-mCherry-expressing rats, CNO infusions had no effect on the density of c-Fos+ cells in the NAc core compared to control levels (Fig. 5F; vs. mCherry-aCSF, z = - 1.91, *p* = 0.06; vs. mCherry-CNO, z = - 1.66, *p* = 0.11; vs. G_q_ DREADD-aCSF, z = - 1.97, *p* = 0.05). Thus, chemogenetic activation of IL→NAc Shell neurons significantly increases c-Fos expression in the NAc Shell.

**Figure 5.**
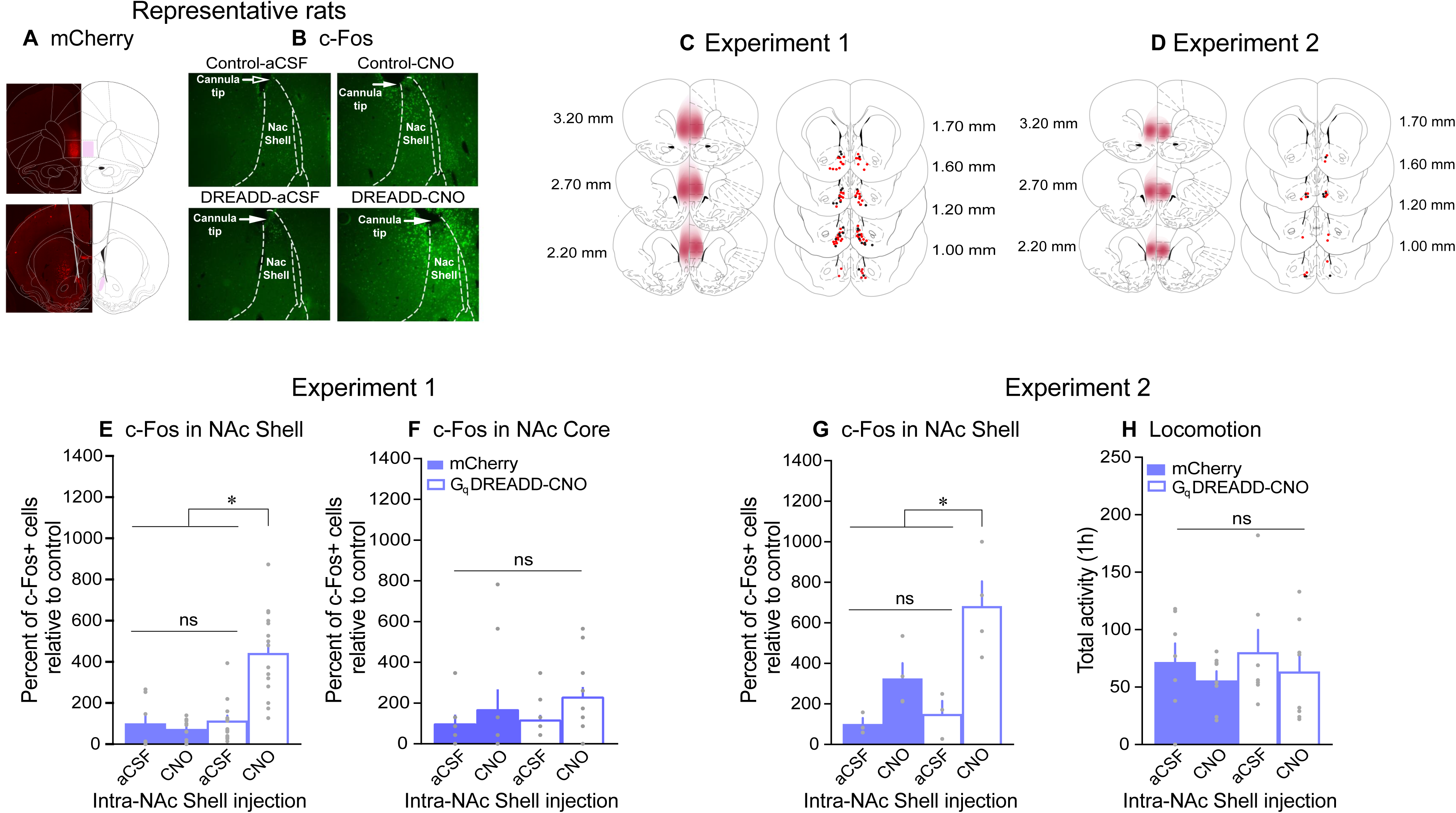
Validation of G_q_ DREADD-mediated activation of IL→NAc Shell neurons and clozapine-N-oxide effects on spontaneous locomotor activity. **A** mCherry fluorescence in the infralimbic cortex (top panel) and estimated injector tract in the nucleus accumbens (NAc) Shell (bottom panel) in a representative rat. **B** c-Fos immunofluorescence in the NAc Shell of representative rats from each group. mCherry expression and representation of estimated injector tip placements in the NAc shell in **C** Experiment 1 and **D** Experiment 2. In Experiment 1, intra-NAc Shell infusions of clozapine-N-oxide (CNO) in rats expressing G_q_ DREADD-mCherry in IL→NAc Shell neurons significantly increased c-Fos protein expression in **E** the NAc Shell but not in **F** the NAc core. In Experiment 1, **G** CNO-mediated chemogenetic activation of IL→NAc Shell neurons significantly increased c-Fos protein expression in the NAc Shell. **H** Intra-NAc Shell CNO injections have no significant effects on spontaneous locomotion. * *p* ≤ 0.01, DREADD-mCherry CNO vs DREADD-mCherry aCSF, mCherry-CNO & mCherry-aCSF. Data are mean ± SEM, Experiment 1 n = 49, Experiment 2 n = 14.

### Experiment 2: Effects of intra-NAc shell CNO injections on locomotion

Because our measure of cue-induced cocaine-seeking behaviour depends on intact motor function, we assessed the extent to which chemogenetic activation of IL→NAc shell neurons influences spontaneous locomotion. Fig. 5H shows the locomotor response to intra-NAc shell aCSF and CNO in rats expressing mCherry or G_q_ DREADD-mCherry. The G_q_ DREADD-mCherry rats that received CNO showed similar rates of locomotion compared to the control groups (vs. mCherry-aCSF, z = - 0.46, *p* = 0.64; vs. mCherry-CNO, z = - 0.47, *p* = 0.64; vs. G_q_ DREADD-aCSF, z = - 0.81, *p* = 0.42). Thus, chemogenetic activation of IL→NAc shell neurons did not significantly influence locomotor behaviour.

## Discussion

DSs are unique and powerful triggers of relapse because they can elicit a conditioned motivational state and drive drug craving and seeking before an individual engages in any seeking behaviour or even consciously experiences desire for drug. The neural circuits mediating this effect have seldom been studied. In rats abstinent from chronic intermittent cocaine intake, we determined how activity in neuronal projections from the IL cortex to the NAc shell contributes to increases in cocaine seeking triggered by drug-associated discriminative and conditioned stimuli. We report four main findings. First, rats learned to discriminate a DS+ from a DS-, such that the DS+ prompted increased cocaine self-administration behaviour and the DS-suppressed responding. Second, after abstinence from intermittent cocaine use, response-independent presentation of the DS+ alone or in combination with the CS+ triggered peaks in cocaine-seeking responses, while the CS+ alone was ineffective (see also [16]). Third, (DS+)-triggered peaks in cocaine-seeking behaviour abated with repeated testing but was reinstated by a priming cocaine injection. Fourth, chemogenetic activation of IL→NAc shell neurons suppressed (DS+)-triggered increases in cocaine seeking with no effects on general motor activity. Thus, following abstinence from intermittent cocaine use, IL inputs to the NAc shell mediate DS-controlled cocaine seeking.

### Discriminative stimulus control of intermittent cocaine intake

With levers present throughout each self-administration session, rats learned to lever press when a DS+ indicated that responding would produce cocaine and to suppress responding when a DS-indicated drug unavailability. During each self-administration session, DS+ periods were immediately followed by DS-periods. Consequently, cocaine concentrations would peak right before each DS-period and begin to decline thereafter [56–58]. When drug concentrations begin to decline below an animals preferred level, the animal normally increases its cocaine intake [59, 60]. In the present study, rats learned to ramp up their cocaine intake during the DS+ and inhibit responding during the DS-, when cocaine seeking would normally be especially pronounced.

Thus, through associative learning, DSs come to control instrumental responding for cocaine under intermittent-access conditions [43, 61, 62].

### Cue-induced cocaine seeking behaviour

We determined the extent to which response-independent presentations of a cocaine-associated DS+ and CS+ increase cocaine-seeking actions in the absence of the drug. Thus, in contrast to most other studies of relapse-like behaviour in animal models, we presented cocaine-associated cues in a manner that neither relied on a drug-seeking response nor reinforced such responses [see also [17, 63–66]. We found that CS+ presentation had no effect on responding compared to that seen during inter-trial intervals when no cues were presented. However, presenting the DS+, alone or in combination with the CS+, triggered increases in lever pressing, even though this was not reinforced by cocaine or any cues. The peaks in responding during DS+ presentation could involve disinhibition of cocaine-seeking due to DS-absence. However, rats did not respond more during inter-trial intervals than they did during DS-presentation. This argues against a disinhibition effect [32]. Instead, our findings support the idea that environmental cues signalling cocaine availability acquire significant incentive properties, enabling them to evoke central conditioned motivational states that in turn prompt increases in relapse-like behaviour [18, 19]. This is reminiscent of findings in humans, where drug cravings are strongest when DSs signal imminent drug availability [67].

Some evidence suggests that the excitatory effects of a DS+ on drug seeking can persist and even incubate across time and repeated testing [15, 68–70], other work shows that these effects begin to extinguish during the very first test session [71]. Consistent with the latter evidence, we found that with repeated testing and/or the passage of time (DS+)-triggered cocaine-seeking behaviour diminished to DS-levels (though DS+ responding levels were still greater than those seen during inter-trial intervals). A priming cocaine injection reinstated cocaine-seeking behaviour specifically during DS+, but not CS+ or DS-presentations (see also [15]). This suggests that being under the effects of cocaine can override previous extinction learning and revive extinguished (DS+)-outcome associations, allowing the DS+ to trigger drug craving and seeking anew. The clinical implication is that extinguishing DS effects might not be enough to reduce the risk of relapse during abstinence, because being in the presence of drug-associated DSs while under the influence of the drug might reinstate DS-controlled drug seeking.

### Contributions of Pavlovian conditioned approach phenotype and prior levels of cocaine intake to cue-induced cocaine-seeking behaviour

Consistent with others [48–50], we found that a sign-vs. goal-tracker phenotype did not significantly predict responding to the DS+ or CS+. Some studies have reported that a goal-tracking phenotype predicts increased (DS+)-guided reward seeking [43, 44], and a sign-tracking phenotype predicts increased (CS+)-guided reward seeking [45]. Notably, some of these previous studies used extinction training before cocaine-seeking tests [43–45], whereas we did not. Previous studies also tested the effects of either DSs or CSs [43, 45], here we tested effects of both cue types in the same rats, in the same test session.

In parallel, we found that rats with the greatest levels of cumulative cocaine intake later responded more to the DS+, CS+ and DS+CS+ combined. Thus, at least some of the inter-subject variability in responding to the cocaine-associated cues can be predicted by the amount of drug previously consumed. Future studies can directly manipulate cocaine intake during the self-administration phase to determine whether this relationship is causal in nature.

### IL→NAc shell projections mediate cocaine seeking triggered by a DS+

Chemogenetic activation of IL→NAc shell suppressed the increases in cocaine seeking triggered by response-independent presentation of a cocaine DS+. By identifying a specific IL-dependent circuit with a causal role in (DS+)-triggered relapse, our findings extend evidence implicating the IL cortex in responding to appetitive DSs [28–35]. Our findings also extend work identifying neural pathways underlying drug-seeking behaviour triggered by contextual DSs (i.e., drug-associated contexts; [20–22, 72]).

Using a CS+, some studies suggest that activity in IL→NAc Shell neurons relies on extinction-induced plasticity to suppress cue-induced cocaine seeking [40]. However other studies also using a CS+ show that extinction is not required [41]. Extinction does not typically occur in human drug users [73, 74], and whether or not there is extinction training can also determine the neurobiological mechanisms underlying cue-controlled relapse [40, 75, 76]. Our rats did not undergo extinction training, and we nevertheless found that activation of IL→NAc shell neurons completely suppressed (DS+)-induced increases in cocaine seeking. Together, our findings and previous work [40, 41, 53] support the idea that rather than mediating extinction-induced plasticity or the associative relationship between specific cue types (i.e., CSs or DSs) and reward, activity in IL→NAc shell neurons guides context-appropriate behavioural responses (e.g., ‘go’ in the presence of drug-associated cues and ‘stop’ following extinction learning [32, 53]) to maximize behavioural utility [77]. Should this be the case, it would make the IL→NAc shell circuit an especially promising target to treat cocaine addiction.

The mechanisms through which IL→NAc shell neurons might modulate (DS+)-induced cocaine seeking remain to be characterized. At the cellular level, cocaine-induced formation of silent synapses might be involved. Cocaine self-administration can generate silent synapses, potentially creating new synaptic contacts, and when these synapses mature/become unsilenced by AMPA receptor insertion, related neurocircuits are remodelled due to strengthened glutamatergic neurotransmission [78–81]. Following extended abstinence (with no extinction training), as done here, cocaine-generated IL→NAc shell silent synapses mature via synaptic insertion of calcium-permeable AMPA receptors [80]. Reversing the maturation of silent synapses in this pathway augments cue-induced cocaine craving, suggesting that silent synapse-based circuit remodelling may be an endogenous antirelapse neuroadaptation [80]. Thus, we predict that increasing activity of the IL→NAc Shell pathway suppresses (DS+)-triggered peaks in cocaine seeking in part by activating unsilenced synapses within this circuit. Future studies can examine this hypothesis.

### Neuroanatomical specificity

We used viral vector-mediated gene transfer to restrict Gq-DREADD expression to NAc shell-projecting IL neurons and local CNO infusions to restrict pharmacological effects to the NAc shell. IL cortex fibres primarily target the caudo-medial portion of the NAc shell, whereas prelimbic cortex fibres are distributed throughout the NAc, but terminate more heavily in the core vs. shell [82–85]. We saw some virus expression in the ventral aspect of the prelimbic cortex. However, CNO injection into the NAc shell did not change the density of c-Fos+ cells the NAc core, suggesting no chemogenetic activation of cells in this region.

### Conclusions

DSs that signal drug availability act as powerful triggers of drug-seeking actions, before any such actions have been initiated. Despite the unique contributions of DSs to relapse vulnerability, few studies have examined neural circuit mechanisms underlying DS effects on drug-seeking behaviour, and these have focused on contextual DSs [20–22, 72]. Here we identify the IL→NAc shell pathway as a key circuit mechanism for pre-empting cue-driven drug seeking—shedding light on mechanisms of relapse before overt behaviour is initiated. To the extent that our procedures in rats are relevant to addiction in humans, our findings suggest that this neural pathway could be an effective target for interventions designed to reduce relapse to cocaine during abstinence.

## Funding and disclosure

Funding This research was supported by a grant from the Canadian institutes of Health Research (grant number 157572) to ANS. HEA was supported by a PhD scholarship from the Libyan cultural Office (LCAO).

## Acknowledgements

We thank Dr. Stephen Mahler for technical advice on implementing chemogenetic manipulations. We also thank Dr. Mike Robinson and Dr. Mandy Lecocq for technical support and advice on data analysis. We are grateful to Dr. Marina Wolf for her comments on this manuscript before journal submission.

## Author contributions

HEA and ANS designed the research. HEA, IL, DC, SN, AP and TP performed experiments. HEA and IL analysed the data with guidance from ANS. HEA and ANS wrote the article. All authors critically reviewed the content and approved the final version for publication

## Declarations

Competing interests: The authors have nothing to disclose.

## Notes

### Competing Interest Statement

The authors have declared no competing interest.

