## Supplementary Methods for "Activation of Infralimbic cortex neurons projecting to the nucleus accumbens shell suppresses discriminative stimulus-triggered relapse to cocaine seeking in rats"

**Subjects**

For Experiment 1, we used 56 female Sprague-Dawley rats (225-250 g; Envigo, Frederick barrier 208A, Maryland, USA). For Experiment 2, we used 14 female Sprague-Dawley rats (150–175 g; Charles River, California, USA). Rats were single-housed in a temperature-controlled room (21°C) on a reverse 12/12 h dark/light cycle (lights off at 8h30 AM) with free access to water and restricted food access. Rats had three days of acclimation to the vivarium before any experimental manipulations. Food (Charles River Purina chow #5075) was then restricted to 18 g/day [1]. Mild food restriction produces healthier rats compared to ad libitum feeding [2, 3] and reduces the mg/kg amount of drug needed during self-administration sessions. All experimental procedures were conducted during the dark phase of the rats' circadian cycle.

**Drugs**

Cocaine hydrochloride (Galenova, St-Hyacinthe, QC, Canada) was dissolved in 0.9% saline and filtered with corning bottle-top filters (0.22 µm PES membrane; Fisher Scientific, Whitby, ON, Canada). Every three days, the cocaine concentration was adjusted according to average rat weight. Clozapine-N-oxide (CNO; Adooq Bioscience, cat#A15048, lot L15048B002, Irvine, CA) was dissolved in artificial cerebrospinal fluid/0.1% dimethyl sulfoxide solution for a final concentration of 1mM.

**Apparatus**

In Experiment 1, All training and testing took place in standard operant cages (Med Associates, St Albans, VT) located in a testing room separate from the rats' housing room. Each cage was equipped with a ventilating fan and two retractable levers. A food port was located between the levers and a white cue light was located above each lever. Another white cue light was located on the back wall of the cage. Pressing the active lever (counterbalanced across the left- and right-side levers) produced reinforcement (sucrose pellet or intravenous cocaine). Pressing the inactive lever had no programmed consequences. Each cage also contained four horizontally aligned photocell beams to measure locomotion during each self-administration session. Locomotion was computed as photocell beam breaks/min.

In Experiment 2, psychomotor activity was measured in Plexiglas cages (27 × 48 × 20 cm) equipped with 6 rows of photocells (3 cm above the floor of the box). These cages automatically recorded photocell breaks as a measure of horizontal locomotor activity.

**Magazine Training**

In preparation for subsequent autoshaping, rats had two magazine training sessions (30 min) to learn to retrieve experimenter-delivered sucrose pellets (45-mg, banana flavoured; Bioserv; product # 76285-260, VWR, Mont-Royal, QC, Canada) from the magazine. The session began with the fan turning on, followed by delivery of 40 sucrose pellets under a variable interval 45 sec schedule (VI45s), without any explicit cues. Levers remained retracted during the session.

**Autoshaping**

Rats first received six 30-min autoshaping sessions (2 session/day). Sessions began with the fan turning on and a lever extending for 8 s (the CS), followed by sucrose pellet delivery (the unconditioned stimulus) under VI45s. Rats received 40 sucrose pellets per session. A response bias score was calculated to identify the rats' phenotype [(CS lever contacts - CS port entries)/(CS lever contacts + CS port entries)]. A score ≤ -0.50 indicates goal-tracking, a score of ≥ +0.50 indicates sign-tracking, and a score of -0.49 to +0.49 indicates an intermediate phenotype [4, 5].

**Sucrose self-administration**

Following autoshaping, rats were trained to press one of the two levers in the operant cage for sucrose pellets under a fixed ratio 1 schedule of reinforcement (FR1), in daily 30-min sessions. The active lever was counterbalanced across the left-side and right-side levers of the cage. Once rats earned ≥ ~ 20 pellets/session on two consecutive sessions, the schedule of reinforcement was changed to FR3 for at least 4 sessions. We presented no reward-associated cues during this phase of training. All rats met the acquisition criterion and were prepared for intravenous catheter implantation.

**Intravenous catheter implantation**

The day following the last sucrose self-administration training session, the rats were anesthetized with isoflurane (5% for induction, 2–3% for maintenance) and catheters were implanted into the right jugular vein, as in previous work [6, 7]. Immediately prior to surgery, rats received penicillin (Derapen, 0.02 ml of 300 mg/ml, i.m.; CDMV, Saint-Hyacinthe, QC, Canada) and an anti-inflammatory agent (Rimadyl, 0.03 ml of a 50 mg/ml solution, s.c.; CDMV, QC, Canada). Thereafter, catheters were flushed on alternate days with saline and saline containing 0.2 mg/ml of heparin (Sigma- Aldrich, Oakville, ON), and 2 mg/ml Baytril (CDMV, St Hyacinthe, QC).

**Cocaine self-administration training**

Following recovery from surgery (7-10 days), rats learned to self-administer i.v. cocaine (0.5 mg/kg/infusion, delivered over 5 s) under FR3, during 1-h sessions, 2 sessions/day. The cage ventilation fan turned on at the beginning of each session. Two min later the levers were inserted and a DS+ (cue light above the left lever) was presented for the remainder of the 1-h session. Under FR3, active lever presses produced a cocaine infusion paired with a 5-s CS+ (cue light above the right lever and sounding of a 2900-Hz, 75-dB tone). Acquisition criteria included taking a minimum of 6 infusions/session and pressing at least twice more on the active versus inactive lever, on two consecutive sessions.

**Intermittent-Access discrimination training**

Rats received 12 intermittent-access (IntA) sessions during which cocaine (0.5 mg/kg/inf, injected over 5 s) was available under FR3 during 5-min ON periods signalled by a DS+ (cue light above the left lever) and then unavailable during 25-min OFF periods signalled by a DS- (cue light on the back wall) [8]. During DS+ periods, each cocaine infusion was also paired with the same 5-s CS+ presented during cocaine self-administration training. The ventilation fan turned on at the beginning of each session. Two min later, the levers were inserted into the cage and the session started with a DS+/cocaine ON period, followed by a DS-/cocaine OFF period. This cycle was repeated 8 times, resulting in a 4-h session (Fig. 2B). Discrimination ratios were computed to determine the ratio of active lever presses during DS+ presentation (total DS+ active lever presses/total DS+ and DS- active lever presses) versus DS- presentation (total DS- active lever presses/total DS+ and DS- active lever presses).

**Catheter patency**

Catheter patency was verified after the final IntA cocaine self-administration session. Rats received an i.v. infusion of the short-acting anesthetic, propofol (10 mg/mL; 0.1 mL/rat; Fresenius Kabi Canada Ltd, Richmond Hill, ON) followed by 0.1 ml saline. Rats that did not become ataxic within 10 s of propofol administration immediately received a 2$^{nd}$ dose. Rats that did not become ataxic after the 2$^{nd}$ dose were considered to have non-patent catheters and were excluded from data analysis.

**Stereotaxic surgery**

Rats first received penicillin (Derapen, 0.02 ml of 300 mg/ml, i.m.; CDMV, Saint-Hyacinthe, QC, Canada) and an anti-inflammatory agent (Rimadyl, 0.03 ml of a 50 mg/ml solution, s.c.; CDMV, QC, Canada). Each rat then received bilateral injections of a viral vector into the IL (A/P + 2.9 and M/L +/- 0.6, in mm relative to Bregma. D/V – 4.8 mm from skull surface above the IL) and had bilateral stainless steel guide cannula (23 gauge; HRS Scientific, Anjou, QC, Canada) implanted into the NAc Shell (A/P + 1.4 and M/L +/- 2.2 at a 10º angle, in mm relative to Bregma. D/V – 5.5 mm from skull surface above the NAc Shell). Microinjector tips extended 1.5 mm beyond cannulae tips for a final DV measurement of -7 mm. In Experiment 1, 36 rats received a viral vector containing a constitutive $G_q$-coupled designer receptor exclusively activated by a designer drug (AAV8-CaMKIIα-hM3D($G_q$)-mCherry; Canadian Neurophotonics Platform - Viral Vector Core, QC, Canada) and 20 rats received a viral vector containing mCherry alone (AAV8-CaMKIIα-mCherry; Addgene, Cambridge, MA). In Experiment 2, seven rats received AAV5-CaMKIIα-hM3D($G_q$)-mCherry (Addgene) and 7 rats received AAV5-CaMKIIα-mCherry (Addgene). Using a glass pipette (tip diameter, 50 µm) coupled to a Nanoject II (Drummond Scientific, Broomall, PA) we delivered 14 microinjections of 36.8 nl each into each IL, at 10-s intervals and for a total volume of ~0.5 ul/hemisphere. After the microinjections, the glass pipette was left in place for 10 min for diffusion.

**Cue-induced cocaine seeking with chemogenetic activation of IL→NAc shell neurons**

Six weeks following the last IntA session (and 5-6 weeks after adeno-associated virus microinjections), we assessed the effects of activating IL→NAc shell neurons on cue-induced cocaine-seeking behaviour. Within the same test session (50 min), the DS+, CS+, DS- and DS+/CS+ combined were presented independent of the rats' behaviour. The session started with a 2-min no-cue period during which the fan was turned on and the two levers were inserted for the remainder of the session. Each cue type was then presented for 2 min, three times in pseudo randomized order, separated by a 2-min inter-trial interval (ITI). At all times, lever pressing produced no cues and no cocaine. Rats received bilateral CNO (0.5 µl/hemisphere) or aCSF microinfusions into the NAc shell ≤ 10 min before the test, in a between-subjects design.

**Microinfusions**

Microinfusions were administered bilaterally into the NAc shell with a 33-gauge injector connected to a 5-µL Hamilton syringe (VWR, Mont-royal, QC, Canada) set on a motorized syringe pump (Harvard Apparatus, PHD 2000, Saint-Laurent, QC, Canada). Microinfusions were given in a volume of 0.50 µl/hemisphere at a rate of 0.25 µl/min

**Histology and c-Fos immunohistochemistry**

Ninety min after bilateral intra-NAc shell microinjections of 1 mM CNO or aCSF, rats were deeply anesthetized with urethane (1.2 g/kg, i.p.) and transcardially perfused using phosphate-buffered saline followed by 4% paraformaldehyde. Fixed brains were removed from the skull, and post-fixed in 4% paraformaldehyde overnight, then stored in 30% sucrose/phosphate-buffered saline (PBS) for 72 h. Brains were then frozen at -20 °C in a cryostat (Leica CM1850, Leica Bio- systems, IL, United States), sectioned coronally at 40 μm and stored in antifreeze solution at -20 ◦C until processing. For immunohistochemistry, we performed all steps at room temperature with gentle agitation. Free-floating slices were permeabilized with 0.3% Triton X-100/PBS for 15 min. Slices were then incubated for 1h in blocking solution containing 10% Normal Goat Serum (NGS) and 0.1% Triton X-100/PBS. Next, slices were incubated in the same blocking solution as above but now containing 2% NGS and Rabbit anti-c-fos 1/2000 antibody (Cell signaling technology, Danvers, MA) for 24 h. The next day, slices were first washed 3 times with PBS and then incubated in the blocking/2% NGS solution containing 1/500 of a secondary antibody (goat anti-rabbit, Jackson ImmunoResearch Labs, West Grove, PA) for 2 h. Finally, we coverslipped slices with Fluoromount G™ (ThermoFisher, Saint-Laurent, QC, Canada) and examined them under an epifluorescent microscope (Nikon Eclipse E600; Nikon, Tokyo, Japan). We acquired images using Simple PCI software (CImaging Systems, Compix Inc., PA) at 10X. We counted c-fos positive cells automatically using a set threshold for detection with Trainable Weka Segmentation Fiji plugins (ImageJ; National Institutes of Health). Injector placement was verified according to the rat brain atlas of Paxinos & Watson [9]. We excluded data from rats that did not express mCherry in the IL cortex (n = 2) or that had misplaced cannulae in either hemisphere (n = 3).

**Statistical analyses**

All experiments used a mixed design with cue condition as the within-subjects factor (DS+, DS-, CS+, DS+/CS+ and ITI) and treatment condition (CNO or aCSF) or cocaine injection (0 or 10 mg/kg, i.p.) as the between-subjects factors. We used the Shapiro-Wilk tests to assess data normality. We used nonparametric tests to analyze non-normally distributed data; Wilcoxon tests to analyze within-subject effects and Mann-Whitney tests to analyze between-subject effects. Only data on latency to respond to the DS+ (Figs. 3E-F and 4E-F) were normally distributed and we analysed these using two-way repeated-measures ANOVA (Group x Trial number; Trial number as a within subjects variable) with Greenhouse–Geisser correction to adjust for lack of sphericity where appropriate. We used Pearson's correlation coefficient to conduct correlations. We conducted statistical analyses using SPSS (Version 29) and created figures with Graphpad Prism (Version 8; La Jolla, CA). We used an $\alpha$ level of $p < 0.05$ and data in figures are means ± SEM.
