## Supplemental Table 1 for "Activation of Infralimbic cortex neurons projecting to the nucleus accumbens shell suppresses discriminative stimulus-triggered relapse to cocaine seeking in rats"

**Table 1:**
**Correlations between each of Pavlovian conditioned response score and cumulative cocaine intake and active lever pressing during cue-induced cocaine seeking testing.**

| | Cue-induced cocaine seeking behaviour | | | | | |
|---|---|---|---|---|---|---|
| | Active lever pressing during DS+ | | Active lever pressing during CS+ | | Active lever pressing during DS+CS+ | |
| | $r^2$ | *p* | $r^2$ | *p* | $r^2$ | *p* |
| **Pavlovian conditioned response score** | 0.02 | 0.41 | 0.02 | 0.47 | 0.01 | 0.52 |
| **Cumulative intake** | 0.20 | 0.009* | 0.30 | 0.001* | 0.18 | 0.02* |
